# RAREsim2: Flexible simulation of rare variant genetic data using real haplotypes

**DOI:** 10.64898/2026.01.14.694480

**Authors:** Jessica I. Murphy, Ryan Barnard, Megan Null, Audrey E. Hendricks

## Abstract

**Summary:** Realistic simulated data is critical for advancing methodological development and optimizing study design in genetics research. However, many genetic simulation tools are unable to replicate the distribution of rare variants or incorporate key genetic information, such as functional annotations and linkage disequilibrium. RAREsim, an accurate rare-variant simulation algorithm that uses real genetic haplotypes, was developed to address these limitations. Here, we introduce RAREsim2, an update that provides both streamlined software and new functionalities for simulating individual-level differences (e.g., cases vs controls, technological or batch effects) and variant-level differences to represent a variety of causal models.

We demonstrate RAREsim2’s utility with three rare variant association methods (Burden, SKAT, and SKAT-O) across several simulation scenarios: causal variants only in cases, causal variants in both cases and controls, and no causal variants. Type I Error was maintained and the optimal test matched previously known patterns: Burden performed best given a large amount of causal variants with the same direction of effect; SKAT performed best given causal variants with opposite directions of effect. SKAT-O was powerful across all simulation scenarios. We highlight RAREsim2’s capabilities to simulate various genetic ancestries (African, East Asian, Non-Finnish European, and South Asian), gene sizes (~20-80 functional rare variants per gene), strengths of association (20%, 40%, 60% functional variants), and proportions of risk variants (1, 0.75, 0.5). Importantly, real genetic regions can be simulated to include known variant functions and disease associations. Ultimately, RAREsim2 offers additional flexibility and ease in simulating a multitude of realistic genetic scenarios.

**Availability and implementation:** The RAREsim2 python package is available at https://github.com/Hendricks-Research-Team/RAREsim2 and the code for the example demonstration is available at https://github.com/JessMurphy/RAREsim2_demo.

## INTRODUCTION

Simulating data that adequately emulates reality is crucial for study design, power analyses, and method development and evaluation. However, in genetics research, simulating a realistic distribution of rare variants (defined here as variants with a minor allele frequency [MAF] < 1%) while preserving variant level information, such as functional and disease associations, is particularly challenging. In 2022, we developed the simulation method and software RAREsim to address these gaps. RAREsim uses real genetic haplotypes to create simulated data that matches the observed AF distributions and linkage disequilibrium (LD) patterns seen in real data. Thus, RAREsim can capture unique characteristics of specific genetic regions and integrate existing variant features such as functional annotations, constrained regions, and disease loci (Null et al., 2022). Despite these advances, gaps remain including the ability to simulate case/control status or other group-level differences.

Here, we present RAREsim2, which streamlines the software and provides additional functionalities. These new capabilities include simulating different groups of individuals, such as cases and controls, to facilitate power analyses and incorporate technological or batch effects. RAREsim2 also supports simulating distinct groups of variants, allowing researchers to model diverse genetic architectures (e.g., causal variants with opposite directions of effect). Below, we describe the RAREsim2 workflow, provide exemplars of its utility, and benchmark its performance. Ultimately, RAREsim2 enables the simulation of rare variant genetic data across a variety of scenarios using a simple and flexible pipeline that can be tailored to fit users’ specific needs. RAREsim2 is available as a python package on the Python Package Index (PyPI).

## IMPROVEMENTS AND UPDATES

### Workflow for Simulating Different Groups of Individuals with Distinct Variant Structures

RAREsim2 can simulate differences within groups of individuals (e.g., cases and controls, datasets from different sources) based on the general workflow below.

1. **<u>Prepare</u>** input files.
  a. **<u>Simulate</u>** haplotypes at the population-level with an over-abundance of rare variants using Hapgen2 (Su et al., 2011). Simulate all haplotypes together to avoid batch effects and inflated type I error.
  b. **<u>Annotate</u>** the legend file (e.g., functional annotations, risk or protective status).
  c. **<u>Estimate</u>** the expected number of functional 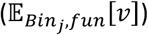 and synonymous 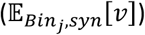 rare variants for each minor allele count (MAC) bin, *j*, based on the allele frequency spectrum (AFS) in the target data and the simulated sample size, using the *calc* function.

i. Use weights *w*_*fun*_ and *w*_*syn*_ (default of 1) to vary the expected number of functional and synonymous variants, respectively, for each group (e.g., *w*_*fun*_ = 1.2 for cases and *w*_*fun*_ = 1 for controls).
2. **<u>Prune</u>** the over-simulated haplotype dataset to match the expected distributions of functional 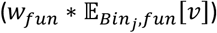 and synonymous 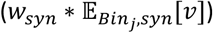 variants estimated from summary-level target data using the *sim* function.
3. **<u>Extract</u>** subsets of 2*N* haplotypes, where *N* is the simulated sample size (e.g., 2*N*_*case*_ or 2*N*_*control*_) using the *extract* function.

Steps 1a and 1b are performed prior to using the RAREsim2 *calc, sim*, and *extract* functions in Steps 1c, 2, and 3 respectively. Steps 2-3 can be repeated to achieve the desired study design. A flowchart of how RAREsim2 can be used to simulate case-control data is in **Figure 1A**, with more detailed versions in the supplemental (Error! Reference source not found.**-**Error! Reference source not found.). The flowchart shows the over-simulated haplotypes, annotated legend file, and expected number of variants are all user inputs for the pruning process (Step 2), which outputs pruned haplotypes and an updated legend file. Random subsets of the pruned haplotypes can then be extracted and designated as cases or controls (Step 3) and/or used as input for additional pruning steps. In addition to case and control groups, other groups of individuals can be simulated, such as external common controls or batch effects (Error! Reference source not found.) (Wojcik et al., 2022).

**Figure 1.**
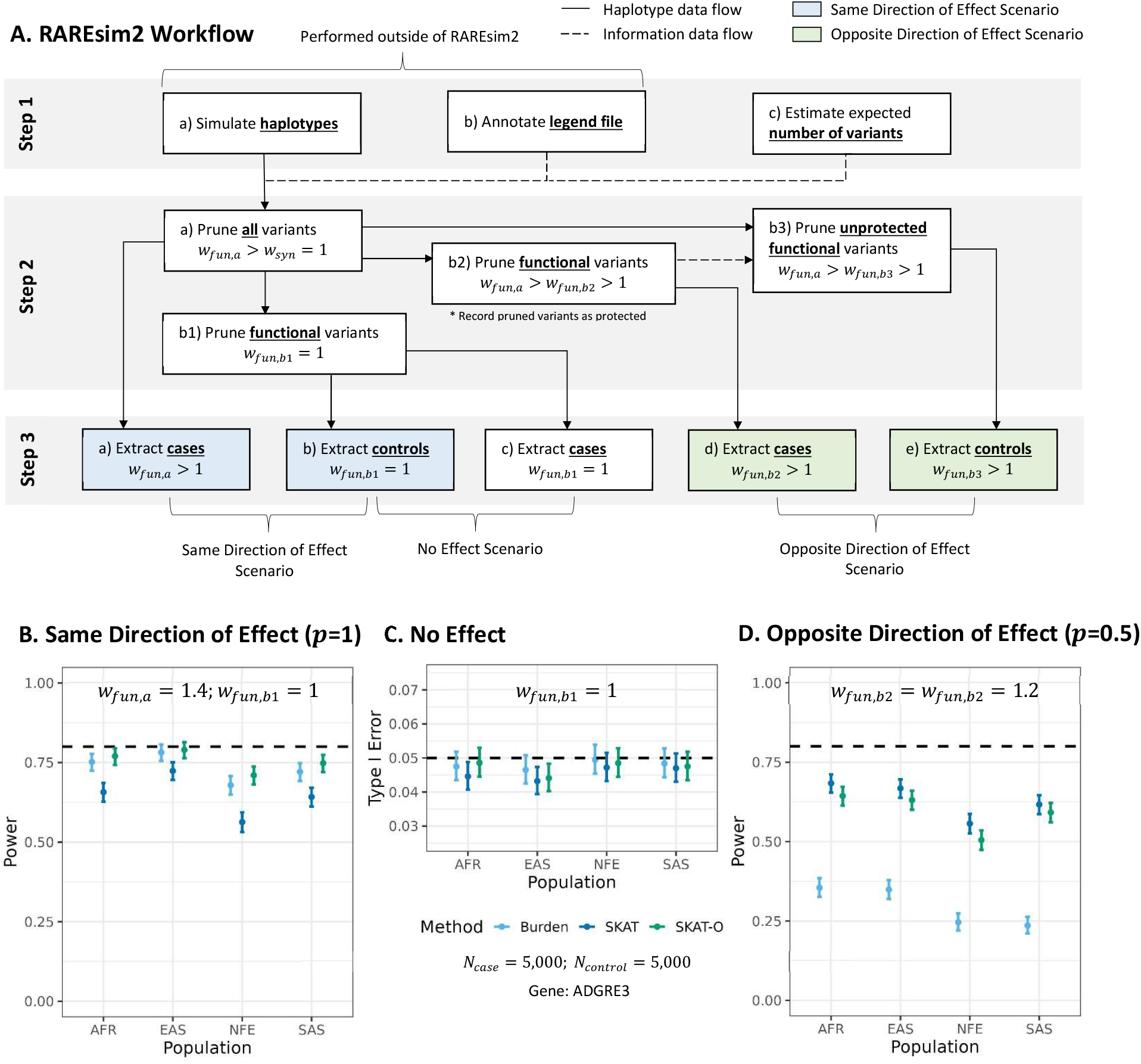
RAREsim2’s general workflow and benchmarking. **(A) General Workflow.** Flowchart for simulating datasets for three scenarios: same direction of effect (i.e., Power, causal risk variants only in cases), no effect (i.e., Type I error, no causal variants), and opposite direction of effect (i.e., Power, causal risk variants in cases and causal protective variants in controls). **(B-D) Benchmarking**. Cases (N=5,000) and controls (N=5,000) were simulated for each scenario using 1000 Genomes reference data and gnomAD target data from four genetic ancestry populations: African (AFR), East Asian (EAS), Non-Finnish European (NFE), and South Asian (SAS). Datasets were created according to the flowchart with weights *w*_*fun,a*_ = 1.4, *w*_*fun,b*1_ = 1, and *w*_*fun,b*2_ = *w*_*fun,b*3_ = 1.2 for Step 2. Results are shown for the *ADGRE3* gene. **(B) Same direction of effect scenario**. As expected, Burden is more powerful than SKAT. **(C) No Effect Scenario**. Type I error is well-controlled (i.e., 0.05) across the Burden, SKAT, and SKAT-O rare variant association tests. **(D) Opposite direction of effect scenario**. As expected, SKAT is much more powerful than Burden. SKAT-O is powerful across the two power scenarios.

### Software Enhancements and Capabilities

RAREsim2’s simulation workflow has been streamlined, enabling users to implement the entire RAREsim2 pipeline in a single program following haplotype simulation in HAPGEN2. Step 1c, which estimates the expected number of variants, can now be performed directly in Python rather than R (as in RAREsim v1). The *calc* function (Step 1c) can accept population default parameters, user-specified inputs, or target data for one or more variant groups. To generate different groups of individuals, users can employ the *extract* function (Step 3) to obtain random subsets of haplotypes, eliminating the need for additional command-line functions. A user-specified random seed can be used to ensure reproducibility and consistent haplotype extraction across differently pruned datasets.

RAREsim2 also introduces a range of new functionalities. Users can now simulate a variety of causal and association models based on variant characteristics designated in the legend file, including functional annotations, pruning probability, or risk or protective status. For instance, pruning can be restricted to functional variants, or protective variants can be excluded from pruning to create groups with more protective causal variants. Additional user control over the number of rare variants is also available with weights to adjust the expected number of total, functional, or synonymous variants, and thresholds to ensure a minimum number of variants observed within each MAC bin.

A complete list of RAREsim2’s new functionalities can be found in Error! Reference source not found..

## RARESIM2 FRAMEWORK FOR CASE-CONTROL SCENARIOS

We used RAREsim2 to simulate three causal variant scenarios (**Figure 1A**): 1) causal risk variants only present in cases (i.e., Power, same direction of effect); 2) no causal variants (i.e., Type I Error); and 3) causal risk variants in cases and causal protective variants in controls (i.e., Power, opposite direction of effect). In **Figure 1A**, we show the steps for each scenario, which are described further below.

For the same direction of effect scenario, we first pruned the haplotypes to have more functional variants than expected (*w*_*fun,a*_ > 1) and approximately the same number of synonymous variants as expected (*w*_*syn*_ = 1; Step 2a in **Figure 1A**). Next, we extracted a random subset of 2*N*_*case*_ haplotypes (Step 3a in **Figure 1A**). Finally, we pruned the functional variants in the haplotypes to approximate their expected distribution (*w*_*fun,b*1_ = 1; Step 2b1 in **Figure 1A**) and extracted a different subset of 2*N*_*control*_ haplotypes (Step 3b in **Figure 1A**). This resulted in cases having more rare functional variants compared to controls (Error! Reference source not found.).

For the no effect scenario (i.e. type I error), we extracted 2*N*_*case*_ haplotypes (Step 3c in **Figure 1A**) using the same haplotype indices from Step 3a and paired them with the internal controls from Step 3b. Adjusting for sample size, the expected number of functional and synonymous variants will be the same for cases and controls (Error! Reference source not found.).

For the opposite direction of effect scenario, we pruned the functional variants from Step 2a to have more variants than expected but fewer variants than in Step 2a (*w*_*fun,a*_ > *w*_*fun,b*2_ > 1; Step 2b2 in **Figure 1A**); the pruned variants were labeled as “protective” in the legend file. From Step 2b2, we extracted 2*N*_*case*_ haplotypes using the same haplotype indices from Step 3a (Step 3d in **Figure 1A**). Then, after excluding the protective functional variants from the haplotypes in Step 2a, we pruned again (*w*_*fun,a*_ > *w*_*fun,b*3_ > 1; Step 2b3 in **Figure 1A**) and extracted 2*N*_*control*_ haplotypes using the haplotype indices from Step 3b (Step 3e in **Figure 1A**). Note, variants pruned from the controls are considered “risk” variants because they are present in the cases but not the controls, whereas variants pruned from the cases but not the controls are considered “protective” because they are in the controls but not the cases. Cases in Step 3d and controls in Step 3e will have the same expected number of functional and synonymous variants after adjusting for sample size if *w*_*fun,b*2_ = *w*_*fun,b*3_, although the actual variants will differ (Error! Reference source not found.).

To ensure the same approximate number of causal and non-causal variants in both power scenarios, the functional weights for the opposite direction of effect scenario (*w*_*fun,b*2_, *w*_*fun,b*3_) were calculated as a function of the weights for Step 2a (*w*_*fun,a*_) and Step 2b1 (*w*_*fun,b*1_) (**Equation 1**)

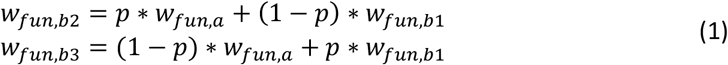

given *w*_*fun,a*_ > *w*_*fun,b*2/*b*3_ > *w*_*fun,b*1_ > *w*_*fun,c*,_ *w*_*fun,b*1_ = *w*_*syn*_ and *w*_*fun,c*_ = *w*_*syn,c*,_ where *p* is the proportion of risk variants and 1 − *p* is the proportion of protective variants with 0 < *p* <1. For example, if we assume 20% of the expected number of functional variants are causal (*w*_*fun,a*_ = 1.2, *w*_*fun,b*1_ = 1) and the proportions of risk and protective variants are equal (*p* = 0.5), then the weights for the case and control sets for the opposite direction of effect scenario would be equal (*w*_*fun,b*2_ = *w*_*fun,b*3_ = 1.1). If the proportions of risk and protective variants are unequal (*p* = 0.75), then the weights would be unequal (*w*_*fun,b*2_ = 1.15, *w*_*fun,b*3_ = 1.05). For all scenarios, the total proportion of causal variants (*w*_*fun,a*_) remains the same (0.1 + 0.1 = 0.15 + 0.05 = 0.2). Here, we assumed *w*_*syn*_ = *w*_*fun,b*1_ = 1, but any values can be used as long as the constraints are satisfied.

## BENCHMARKING DEMONSTRATION

### Computing Environment

We used a high-performance computing cluster for the simulations and ran the RAREsim2 workflow using a singularity container with Hagen2 version 2.2.0, python version 3.10.6, and R version 4.2.1. See the RAREsim2_demo Github repository for more details on the nodes and cores of the computing cluster, specific versions of the R packages, and runtimes for each step of the pipeline.

### Input Data and Simulation Parameters

We used reference data from 1000 Genomes phase 3 (hg19) to simulate a centimorgan block on chromosome 19 with the median number of base pairs (19,029) in four ancestral populations: African (AFR), East Asian (EAS), Non-Finnish European (NFE), and South Asian (SAS) (1000 Genomes Project Consortium, 2015). As described in the original RAREsim paper, we modified the reference haplotype and legend files to include information (e.g. genomic position, reference allele, alternate allele) at each sequencing base within the canonical coding region (Null et al., 2022). Using this modified reference data, we simulated haplotypes in Hapgen2 with an over-abundance of rare variants (Step 1a in **Figure 1A**). We annotated the simulated data using the refGene database in ANNOVAR, classifying functional variants as nonsynomous, stopgain, or stoploss variants and synonymous variants as synonymous variants. (Step 1b in **Figure 1A**) (Wang et al., 2010). We then used target summary data from gnomAD v2.1 to calculate the expected number of variants for the simulated sample size (Step 1c in **Figure 1A**) (Karczewski et al., 2020).

We used the weights provided in Error! Reference source not found. to simulate different scenarios (Step 2 in **Figure 1A**) and extracted sample sizes of *N*_*case*_ = *N*_*control*_ = 5,000 for each scenario (Step 3 in **Figure 1A**). We filtered the pruned datasets to exonic functional variants with MAF<1% in the entire sample. The number of rare functional variants in the target dataset per gene per population are shown in Error! Reference source not found.Error! Reference source not found.. We assessed the concordance between the simulated and target AFS distributions in each ancestry for three genes by calculating the difference in proportions of functional variants in each MAC bin.

### Rare Variant Association Methods

We then applied several rare variant association methods to assess RAREsim2’s ability to generate realistic genetic data across a range of causal variant patterns. Two main classes of rare variant tests are burden and variance component tests (Li & Leal, 2008; Wu et al., 2011). Prior work has shown that burden tests are most powerful when there is a large proportion of causal variants with the same direction of effect while variance component tests, such as SKAT, are most powerful when there are causal variants with opposite directions of effect. Optimal tests, which combine both burden and variance component tests, such as SKAT-O, are powerful across most scenarios (Lee et al., 2012).

Here, we benchmarked RAREsim2’s performance by assessing the power and type I error of Burden, SKAT, and SKAT-O when causal variants have the same, opposite, or no effects (Lee et al., 2023). We calculated type I error and power for 10,000 and 1,000 simulation replicates per scenario, respectively.

## Results

Simulations matched the target AFS distributions although singletons were slightly underrepresented for the small gene (**Error! Reference source not found**.-**Error! Reference source not found**.). Type I error of 0.05 was well-controlled across all methods (**Figure 1C**) and for genes of varying sizes (Error! Reference source not found.). As expected, Burden was more powerful than SKAT for the same direction of effect scenarios (**Figure 1B**), whereas SKAT was substantially more powerful for the opposite direction of effect scenarios (**Figure 1D**). SKAT-O maintained high power for all scenarios. These results were consistent for genes of varying sizes, with the power of each test increasing as the number of functional rare variants in a gene increased (Error! Reference source not found.). Overall, Burden required larger proportions of causal variants (*w*_*fun,a*_ > 1.2) before it outperformed SKAT in the same direction of effect scenario, whereas SKAT remained substantially more powerful than Burden across all proportions of causal variants for the opposite direction of effect scenarios. The power of all methods decreased as the proportion of risk variants (*p*) decreased, with the largest decrease observed for Burden. Power increased for all methods as the number of causal variants (*w*_*fun,a*_) increased (Error! Reference source not found.**-**Error! Reference source not found.). All trends matched expectations supporting RAREsim2 simulations were performing as expected.

## CONCLUSION

We present RAREsim2, a streamlined and highly flexible tool for simulating rare genetic variants from real data. The software allows users to simulate a variety of study designs (e.g., case-control, batch effects) and genetic architectures (e.g., different proportions of causal variants, directions of effect, known genetic associations). We benchmarked RAREsim2’s performance using Burden, SKAT, and SKAT-O, demonstrating its capacity to generate realistic datasets. By simulating rare variant data while preserving key genetic features, RAREsim2 enables more accurate modeling of true genomic complexity, supporting improved study design, method development, and evaluation.

## Acknowledgements

This work was supported by the National Human Genome Research Institute (R35HG011293). This work also used the computing resources at the Center for Computational Mathematics, University of Colorado Denver, including the Alderaan cluster, supported by the National Science Foundation award OAC-2019089. None of this research would be possible without publicly available data from gnomAD and the 1000 Genomes Project.

