## Supplemental Information for "RAREsim2: Flexible simulation of rare variant genetic data using real haplotypes"

#### Updated Functionalities

Functions/arguments highlighted in blue in the table below are completely new RAREsim2 functionalities, while the grey highlighted ones are new to python (but were present in the previous R package). Non-highlighted rows are existing functionalities.

**Table S1.** List of updated RAREsim2 functionalities.

| Function | Argument | Description |
|---|---|---|
| calc | | Calculates the number of expected variants per MAC bin using default population parameters, user-provided parameters, or target data. |
| | --mac | MAC bin bounds (lower and upper allele counts) for the simulated sample size |
| | -o | Output file name |
| | -N | Simulated sample size |
| | --reg_size | Size of simulated genetic region in kilobases (Kb) |
| | --pop | Population (AFR, EAS, NFE, or SAS) to use default values for if not providing alpha, beta, omega, phi, and b values or target data |
| | --omega | Scaling parameter to estimate the expected number of variants per (Kb) for sample size N (range of 0-1) |
| | --phi | Shape parameter to estimate the expected number of variants per (Kb) for sample size N (must be > 0) |
| | --alpha | Shape parameter to estimate the expected AFS distribution (must be > 0) |
| | --beta | Shape parameter to estimate the expected AFS distribution |
| | -b | Scale parameter to estimate the expected AFS distribution |
| | --nvar_target_data | Target downsampling data with the number of variants per Kb to estimate the expected number of variants per Kb for sample size N |
| | --afs_target_data | Target AFS data with the proportion of variants per MAC bin to estimate the expected AFS distribution |
| | -w | Weight to multiply the expected number of variants by in non-stratified simulations (default value of 1) |

| | | |
|---|---|---|
| | --w_fun | Weight to multiply the expected number of functional variants by in stratified simulations (default value of 1) |
| | --w_syn | Weight to multiply the expected number of synonymous variants by in stratified simulations (default value of 1) |
| sim | | Simulates new allele frequencies given input haplotypes, legend file, and expected variants. Now outputs a list of pruned variants (.legend-pruned-variants). |
| | -m | Input sparse matrix file |
| | -H | Output compressed haplotype file |
| | -l | Input legend file |
| | -L | Output legend file (only required when using the -z argument) |
| | -b | Expected number of functional and synonymous variants per MAC bin |
| | --functional_bins | Expected number of variants per MAC bin for functional variants. Must be used with the synonymous_bins argument. |
| | --synonymous_bins | Expected number of variants per MAC bin for synonymous variants. Must be used with the functional_bins argument. |
| | --f_only | Expected number of variants per MAC bin for only functional variants. |
| | --s_only | Expected number of variants per MAC bin for only synonymous variants. |
| | -prob | Variants are pruned allele by allele given a probability of removal in the legend file. |
| | --keep_protected | Variants designated with a 1 in the protected column of the legend file will not be pruned. |
| | --stop_threshold | Percentage threshold for stopping the pruning process (0-100). Prevents the number of variants from falling below the specified percentage of the expected count for any given MAC bin during pruning (default value of 20) |
| | --activation_threshold | Percentage threshold for activating the pruning process (0-100). Requires that the actual number of variants for a MAC bin must be more than the given percentage different from the expected number to activate pruning on the bin (default value of 10) |

| | | |
|---|---|---|
| | --small_sample | Allows for simulation of small sample sizes less than 10,000 haplotypes. However, the recommended method to simulate small sample sizes is to oversimulate the number of haplotypes and randomly down-sample to the desired sample size. |
| | -z+ | Monomorphic and pruned variants (i.e., rows of zeros) are removed from the output haplotype file |
| extract | | Randomly extracts a subset of haplotypes (.haps-sample.gz) and outputs the remaining haplotypes separately (.haps-remainder.gz). |
| | -i | Input haplotype file |
| | -o | Output haplotype file name |
| | -n | Number of haplotypes to extract |
| | --seed | Optional seed for reproducibility |

+If the input haplotype file contains monomorphic variants (i.e., initial rows of zeros) when using the z flag, then the pruned-variants file will contain both monomorphic and actual pruned variants.

### RAREsim2 Flowcharts and Parameters

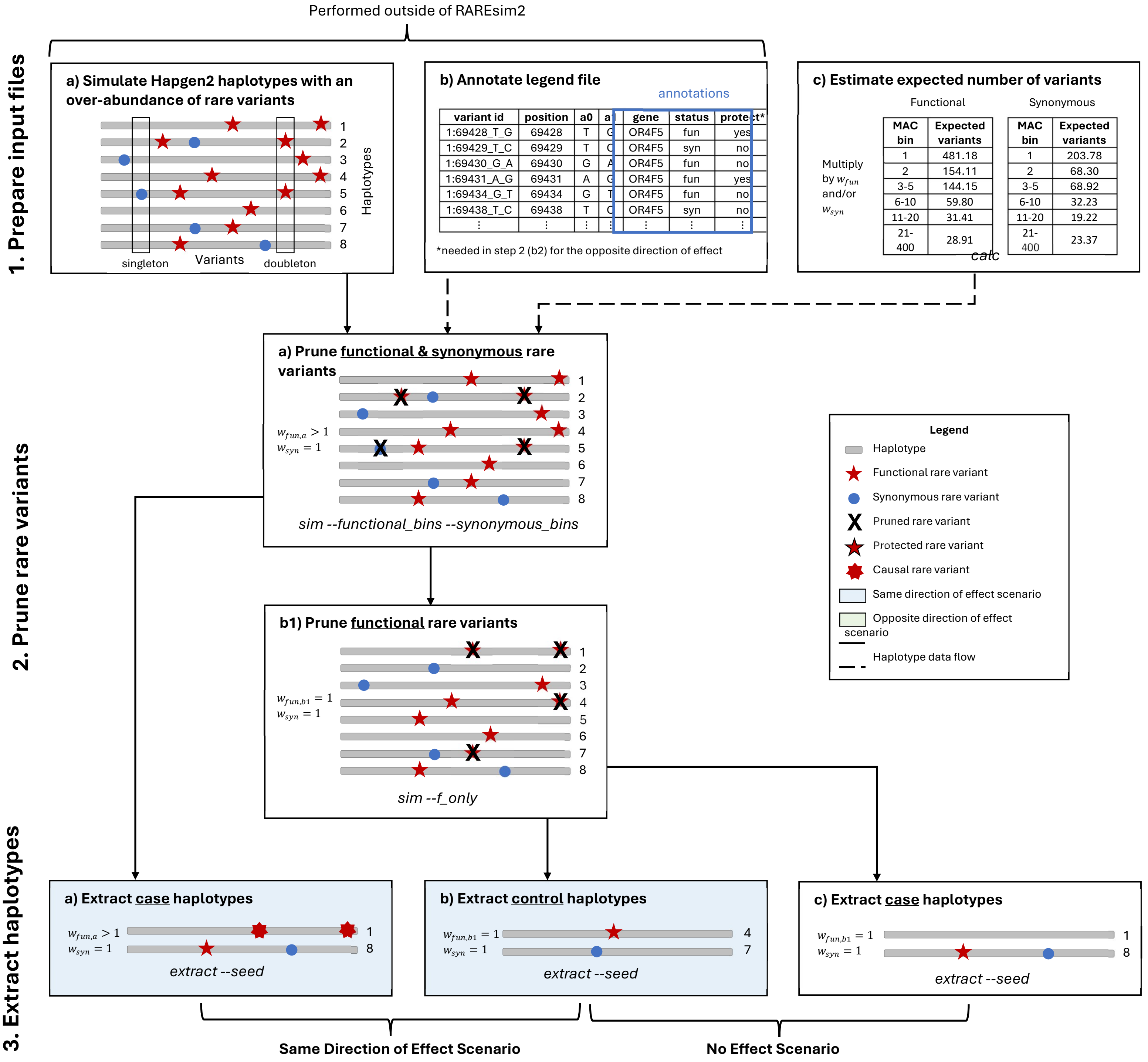

**Figure S1. Detailed RAREsim2 case and control simulation algorithm for the same direction of effect and no effect scenarios**. **| Steps 1a-b.** To simulate an over-abundance of rare variants, the haplotype and legend files are modified to include placeholders for all sequencing bases within coding regions. Eight haplotypes are shown with 12 variants (8 functional in red and 4 synonymous in blue). The initial legend file is amended to include gene and functional status annotations. The legend file can also be updated with the protected status of variants (e.g., for the opposite direction of effect scenario). *Steps 1a* and *1b* are performed prior to RAREsim2. **| Step 1c.** The expected number of variants per MAC bin are estimated for the desired simulation sample size using population default parameters, user-specified parameters, or target data. *Step 1c* may be repeated for different $w_{fun}$ and $w_{syn}$ values. **| Same direction of effect scenario | Step 2a.** The haplotypes are pruned to have more functional variants than expected ($w_{fun,a} > 1$) and approximately the same number of synonymous variants as expected ($w_{syn} = 1$). **| Step 3a.** A random subset of $2N_{case}$ haplotypes are extracted from *Step 2a*. **| Step 2b1.** The functional variants in the haplotypes were pruned to approximate their expected distribution ($w_{fun,b1} = 1$). **| Step 3b.** A different subset of $2N_{control}$ haplotypes are extracted from *Step 2b1*. This results in the cases in *Step 3a* having more rare functional variants compared to the controls in *Step 3b*. **| No effect scenario | Step 3c.** From *Step 2b1*, $2N_{control}$ haplotypes are extracted using the same haplotype indices from *Step 3a* (1 and 8). These cases are paired with the internal controls from *Step 3b* for the no effect scenario. Adjusting for sample size, the expected number of functional and synonymous variants will be the same for cases and controls. The italicized text at the bottom of each box show the relevant functions and arguments needed for each step.

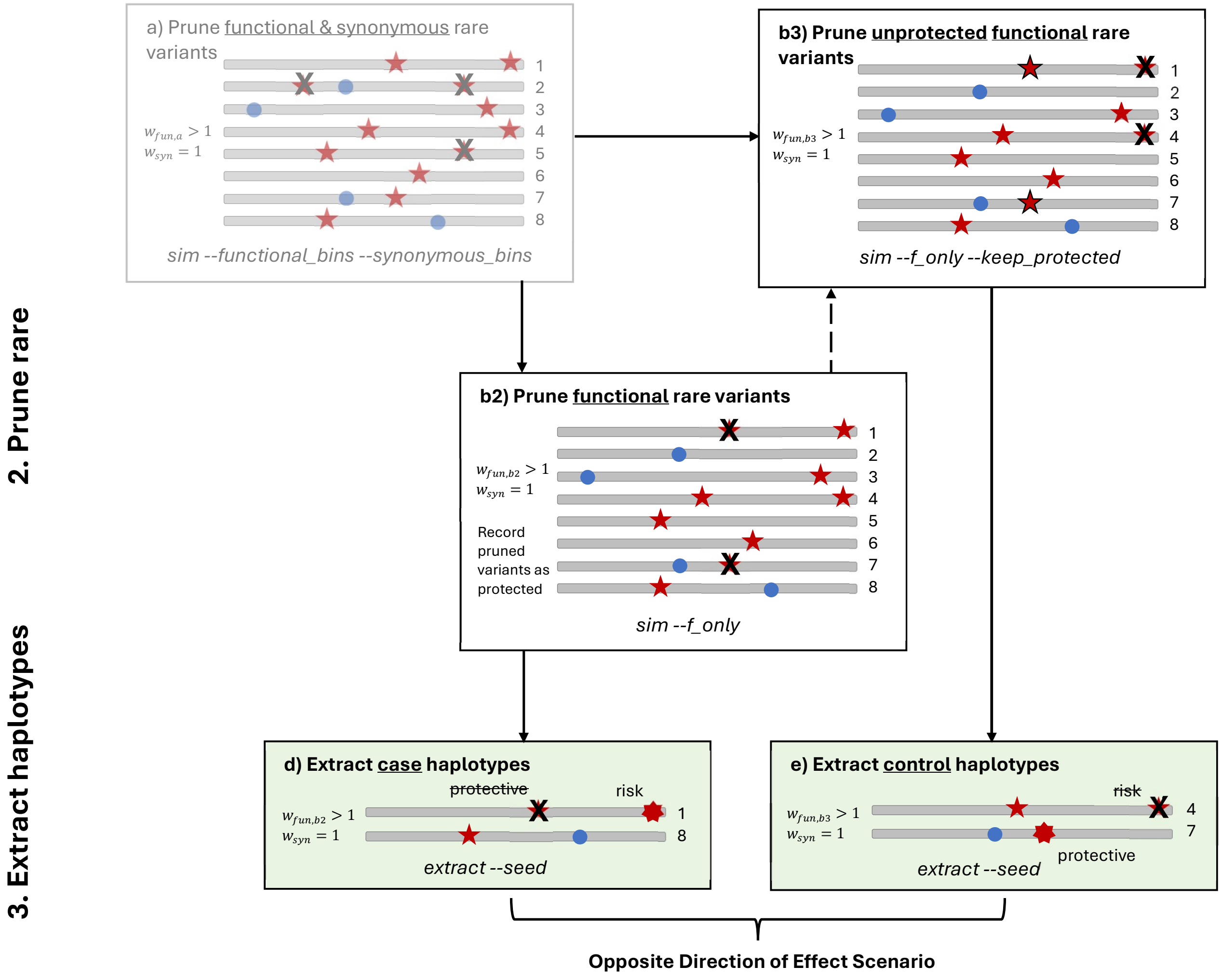


**Figure S2. Detailed RAREsim2 case and control simulation algorithm for the opposite direction of effect scenario.** This is a continuation of Figure S1 *Step 2a*. **| Step 2b2.** The functional variants from *Step 2a* are pruned to less than expected ($w_{fun,a} > w_{fun,b2} > 1$) and the pruned variants are labeled as "protective" in the legend file. **| Step 3d.** From *Step 2b2*, $2N_{case}$ haplotypes are extracted using the same haplotype indices from *Step 3a* and *3c* (1 and 8). **| Step 2b3.** After excluding the protective functional variants from the haplotypes in *Step 2a*, the unprotected functional variants are pruned again ($w_{fun,a} > w_{fun,b3} > 1$). **| Step 3e.** From *Step 2b3*, $2N_{control}$ haplotypes are extracted using the same haplotype indices from *Step 3b* (4 and 7). Note, variants pruned from the controls are considered "risk" variants because they are present in the cases but not the controls, whereas variants pruned from the cases but not the controls are considered "protective". Cases in *Step 3d* and controls in *Step 3e* will have the same expected number of functional and synonymous variants after adjusting for sample size if $w_{fun,b2} = w_{fun,b3}$, although the actual variants will differ.

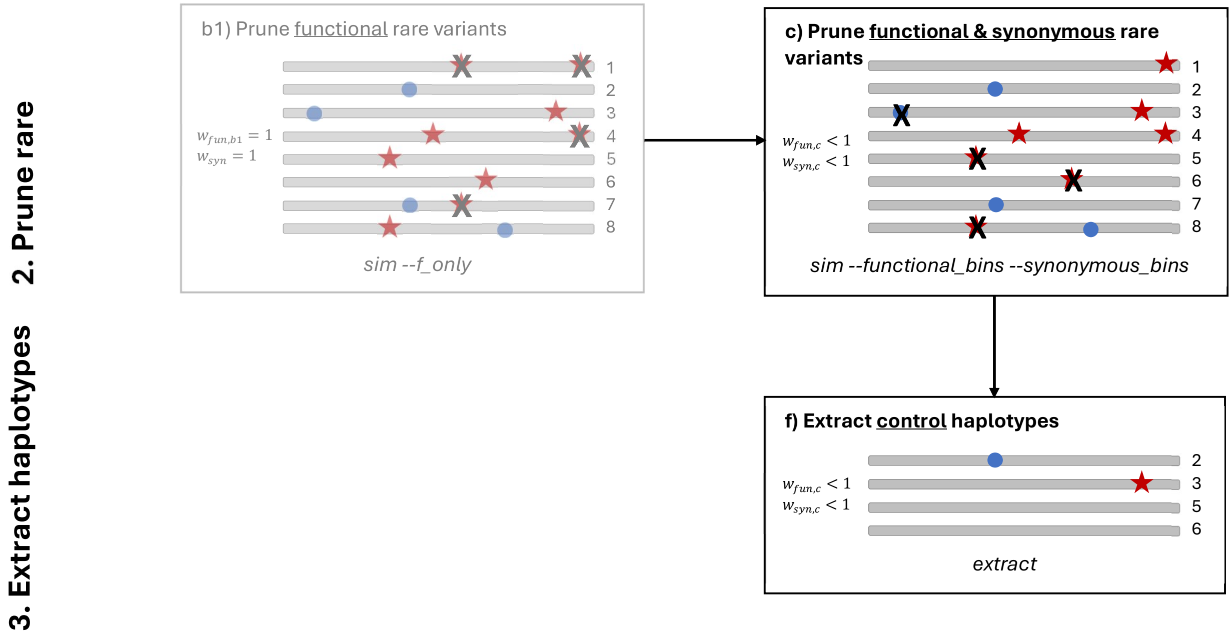


**Figure S3. Detailed RAREsim2 case and control simulation algorithm for confounded external controls.** This is a continuation of Figure S1 *Step 2b1*. **| Step 2c.** The functional and synonymous variants are pruned to less than expected ($w_{fun,c} = w_{syn,c} < 1$). **| Step 3f.** A different subset of haplotypes is extracted as "external controls".

**Table S2.** Functional weights used for generating the datasets for each scenario in the benchmarking example.

| Causal Variant % | Same ($p$=1) | Opposite ($p$=0.75) | | Opposite ($p$=0.5) | |
|---|---|---|---|---|---|
| | $w_{fun,a}$ | $w_{fun,b2}$ | $w_{fun,b3}$ | $w_{fun,b2}$ | $w_{fun,b3}$ |
| **20%** | 1.2 | 1.15 | 1.05 | 1.1 | 1.1 |
| **40%** | 1.4 | 1.3 | 1.1 | 1.2 | 1.2 |
| **60%** | 1.6 | 1.45 | 1.15 | 1.3 | 1.3 |

### Gene Information

**Table S3.** Number of functional rare (MAF<1%) variants in the target dataset (gnomAD v2.1) per gene per population for the median centimorgan block on chromosome 19.

| **Gene*** | **Start** | **End** | **AFR** (N=8,128) | **EAS** (N=9,197) | **NFE** (N=56,885) | **SAS** (N=15,308) |
|---|---|---|---|---|---|---|
| ADGRE5 (large**) | 14491313 | 14519537 | 91 | 78 | 244 | 119 |
| DDX39A | 14519631 | 14530192 | 22 | 22 | 78 | 33 |
| PKN1 | 14543865 | 14582679 | 72 | 69 | 279 | 123 |
| PTGER1 | 14583278 | 14586174 | 21 | 23 | 65 | 39 |
| GIPC1 | 14588572 | 14606944 | 36 | 33 | 113 | 51 |
| DNAJB1 | 14625582 | 14640582 | 29 | 25 | 99 | 46 |
| TECR (small**) | 14627897 | 14676792 | 9 | 18 | 60 | 25 |
| NDUFB7 | 14676890 | 14682874 | 21 | 15 | 45 | 19 |
| CLEC17A | 14693896 | 14721969 | 24 | 24 | 90 | 35 |
| ADGRE3 (medium**) | 14729929 | 14800839 | 72 | 49 | 167 | 70 |
| ZNF333 | 14800613 | 14844558 | 64 | 60 | 182 | 102 |
| ADGRE2 | 14843205 | 14889353 | 102 | 73 | 244 | 131 |

*Highlighted genes are depicted in the Supplemental Results section. The blue highlighted gene is also depicted in Figure 1 of the main paper.
**Sizes refer to the number of variants within the gene - a large, medium, or small amount of rare functional variants compared to the other genes in the centimorgan block.

The figures below compare the AFS distributions in the simulated and target (gnomAD v2.1) data for the three exemplar genes among the four populations. Since the sample sizes differ among the populations in the target data (see **Table S3** above), the proportions of rare functional variants in the simulated (N=10,000) and target data were calculated instead of counts. Then, the difference in proportions between the two were plotted. Smaller boxes centered around 0 (dashed line) represent higher concordance between the simulated and target distributions. As expected, the small gene has much more variability than the medium and large genes. The singletons are also under-estimated in the small gene, whereas they are slightly over-estimated in the large gene.

**Gene: TECR (small)**

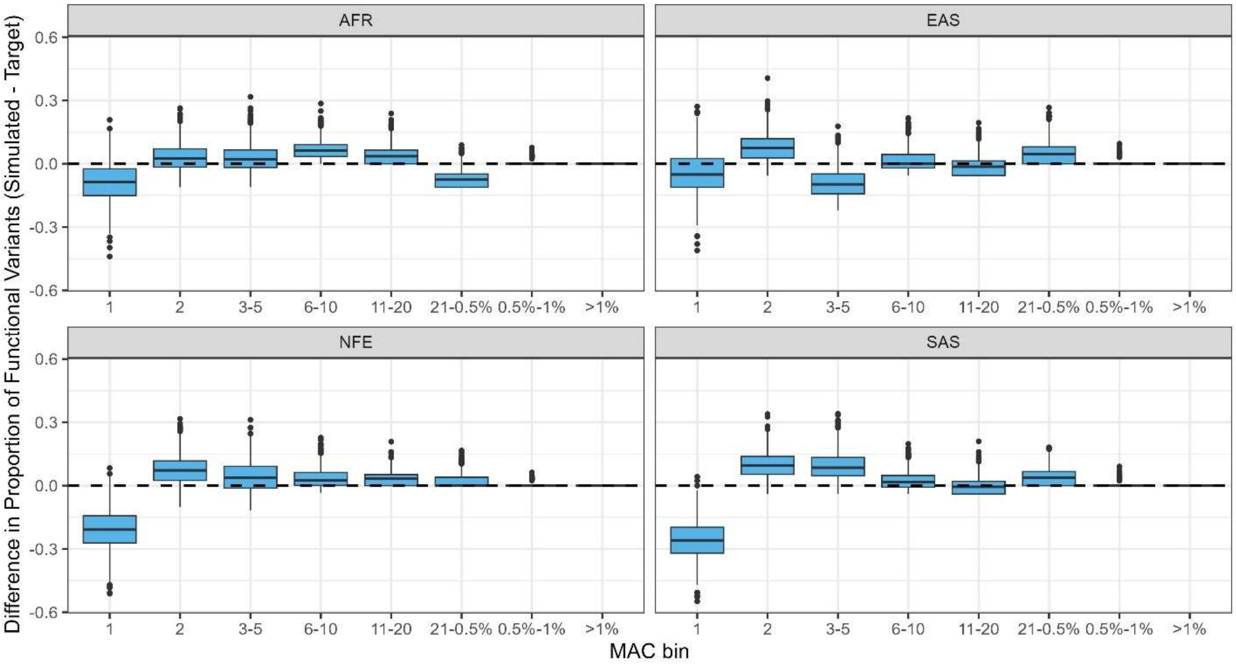


**Figure S4**. Comparison of AFS distributions in the simulated and target (gnomAD v2.1) data for the TECR gene. The percentages in the MAC bin x-axis labels represent MAFs.

**Gene: ADGRE3 (medium)**

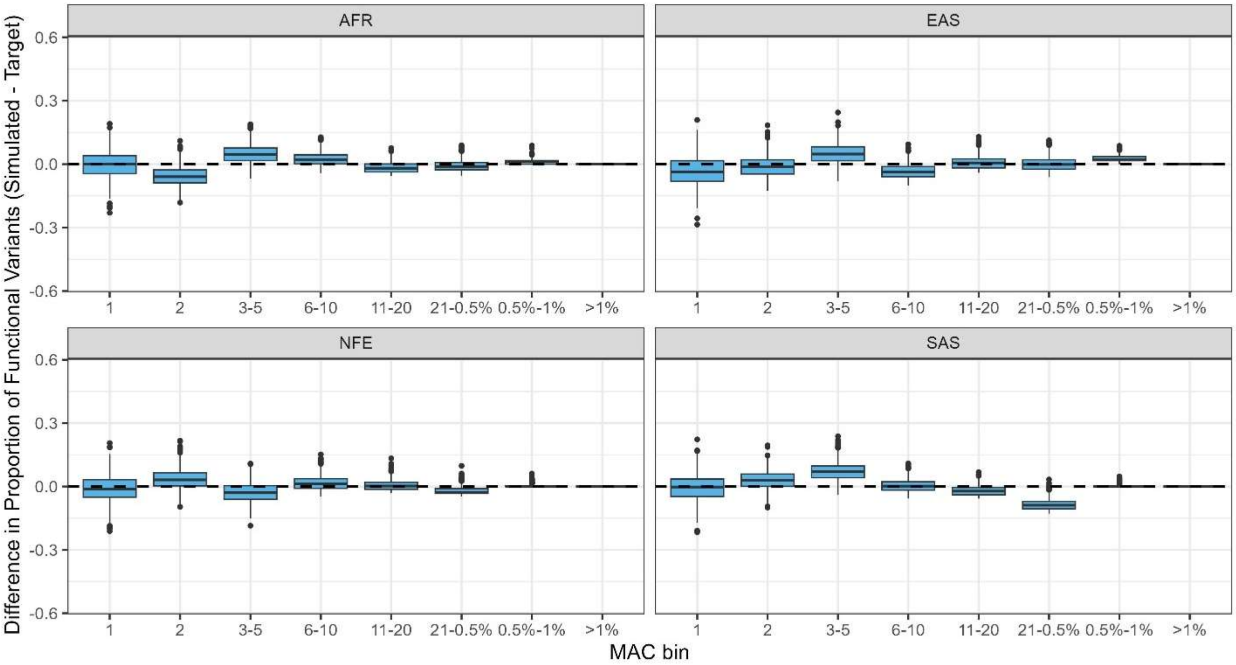


**Figure S5**. Comparison of AFS distributions in the simulated and target (gnomAD v2.1) data for the ADGRE3 gene. The percentages in the MAC bin x-axis labels represent MAFs.

**Gene: ADGRE5 (large)**

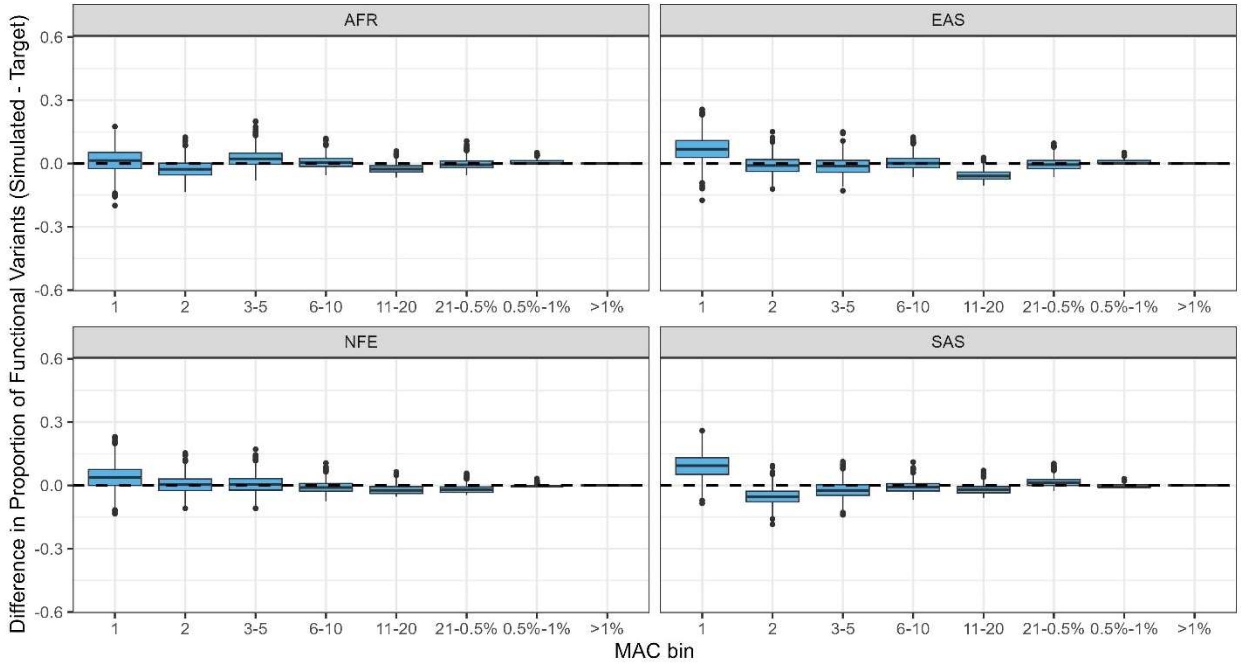


**Figure S6**. Comparison of AFS distributions in the simulated and target (gnomAD v2.1) data for the ADGRE5 gene. The percentages in the MAC bin x-axis labels represent MAFs.

### Results

**No Effect Scenario**

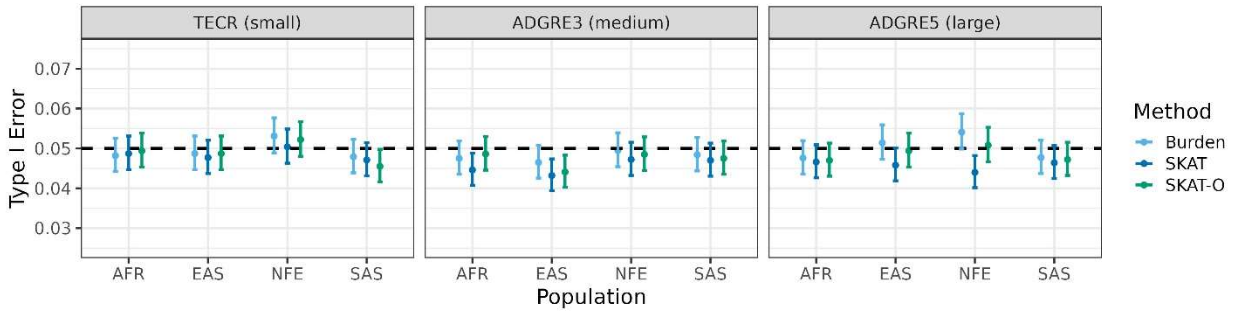


**Figure S7.** No effect scenario (i.e., type I error) results.

**Same Direction of Effect Scenario ($p$=1)**

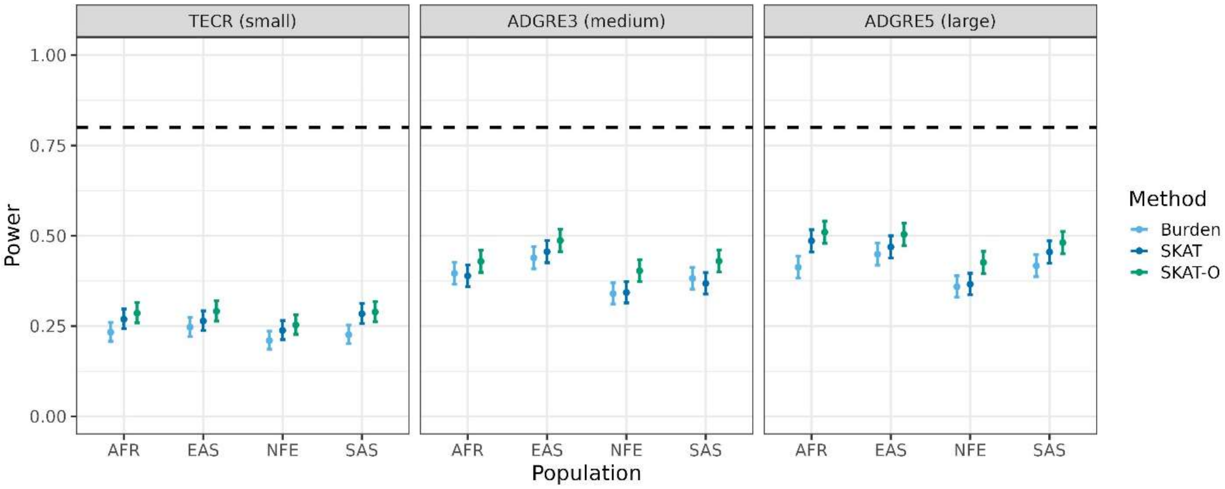


**Opposite Direction of Effect Scenario ($p$=0.75)**

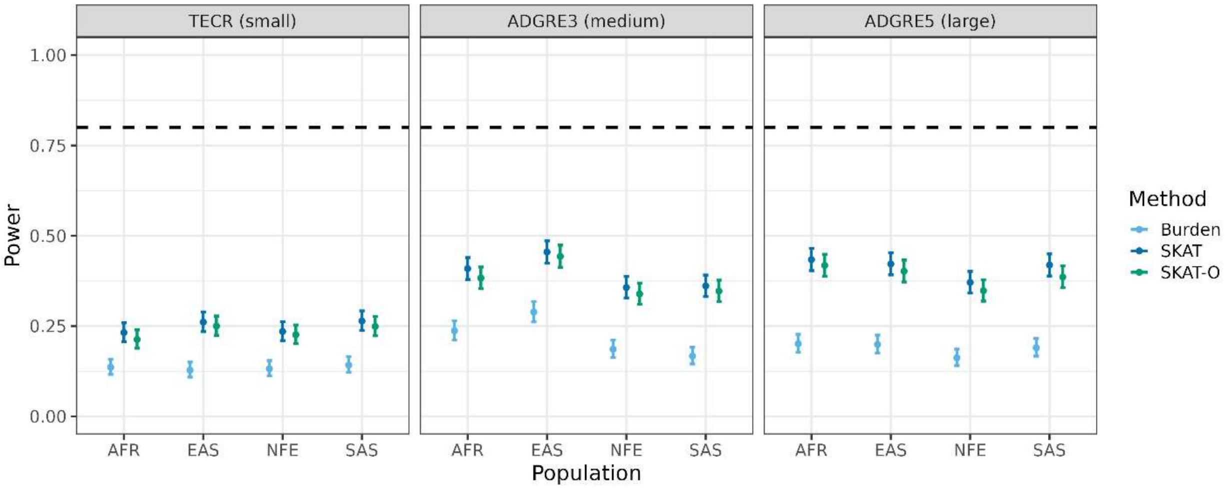


**Opposite Direction of Effect Scenario ($p$=0.5)**

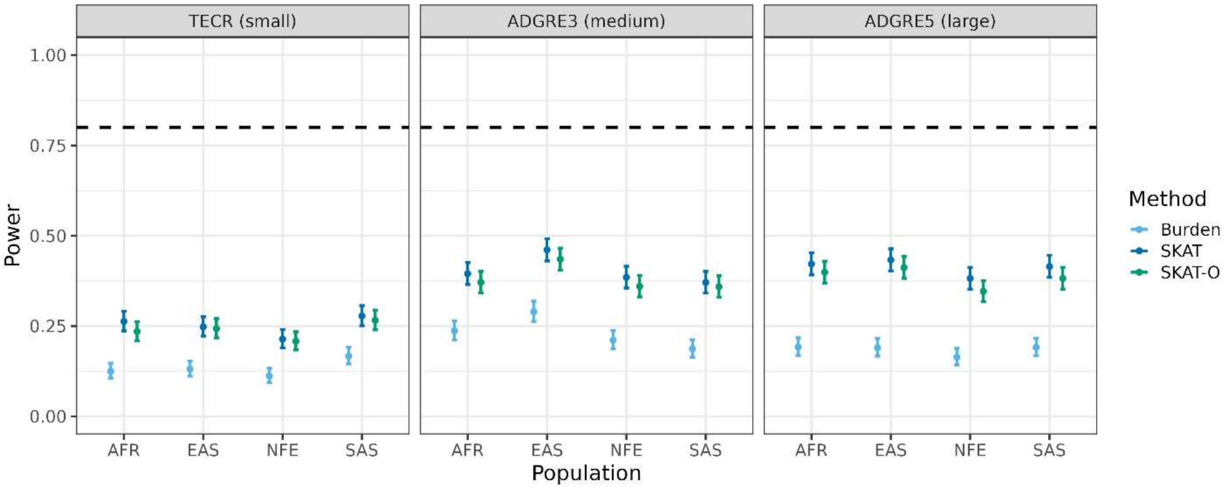


**Figure S8.** Power scenario results for 20% more causal functional variants than expected with $w_{fun,a} = 1.2$, $w_{fun,b} = 1$, and $w_{fun,c1} = w_{fun,c2} = 1.1$.

**Same Direction of Effect Scenario ($p$=1)**

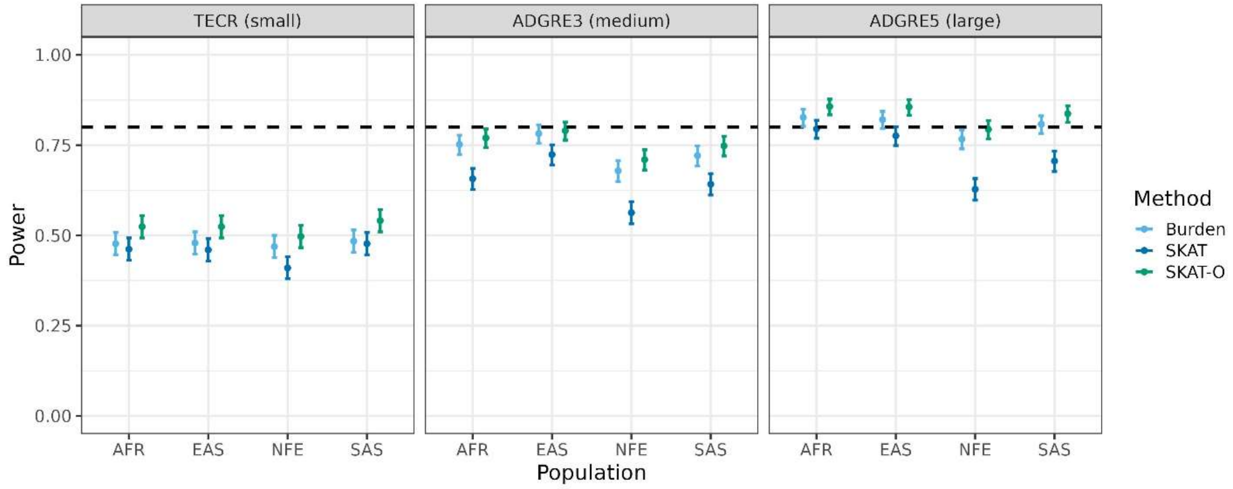


**Opposite Direction of Effect Scenario ($p$=0.75)**

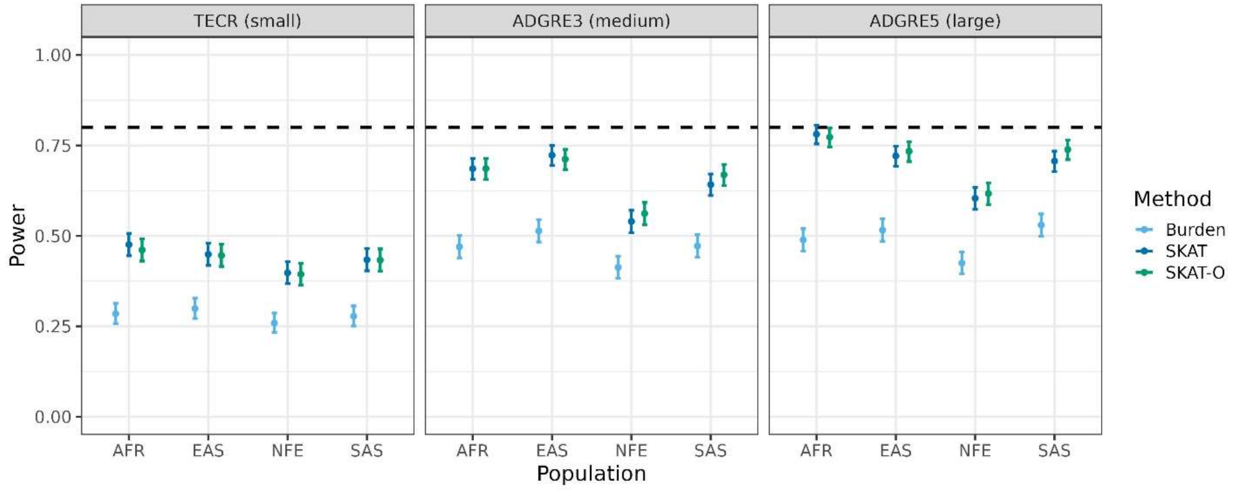


**Opposite Direction of Effect Scenario ($p$=0.5)**

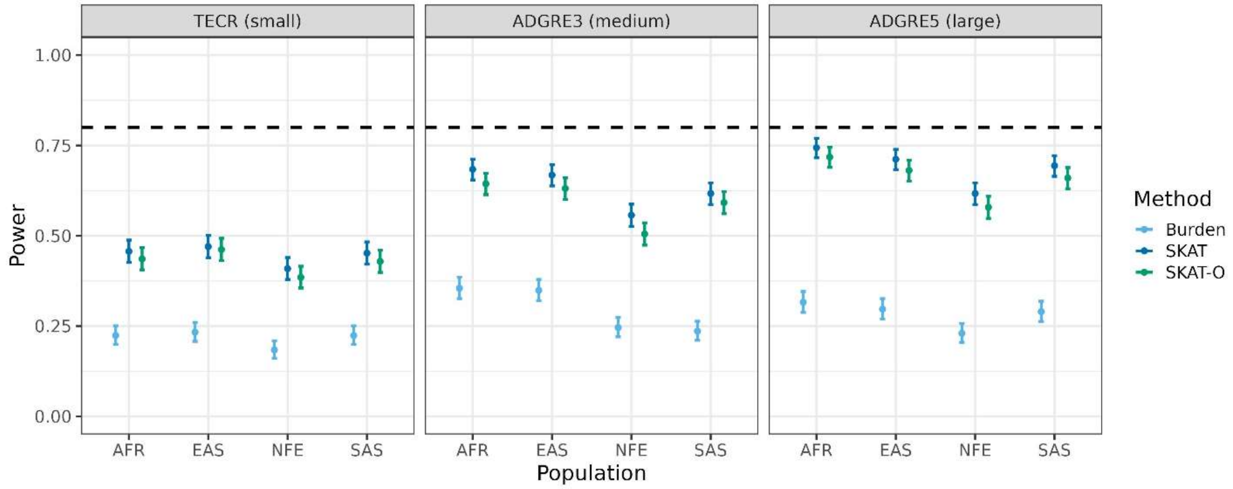


**Figure S9.** Power scenario results for 40% more causal functional variants than expected with $w_{fun,a} = 1.4$, $w_{fun,b} = 1$, and $w_{fun,c1} = w_{fun,c2} = 1.2$.

**Same Direction of Effect *Scenario* ($p$=1)**

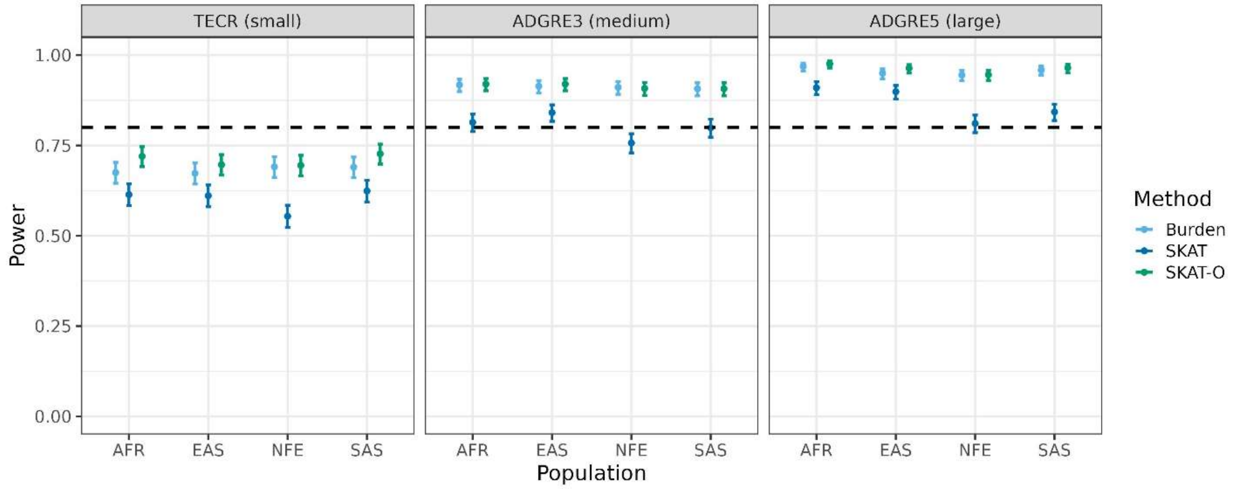


**Opposite Direction of Effect Scenario ($p$=0.75)**

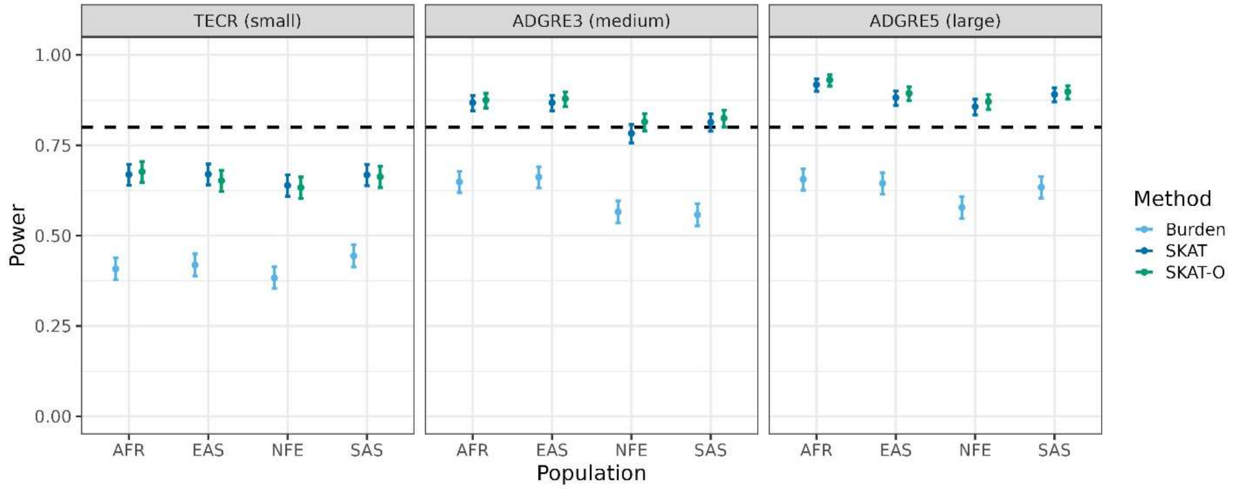


**Opposite Direction of Effect Scenario ($p$=0.5)**

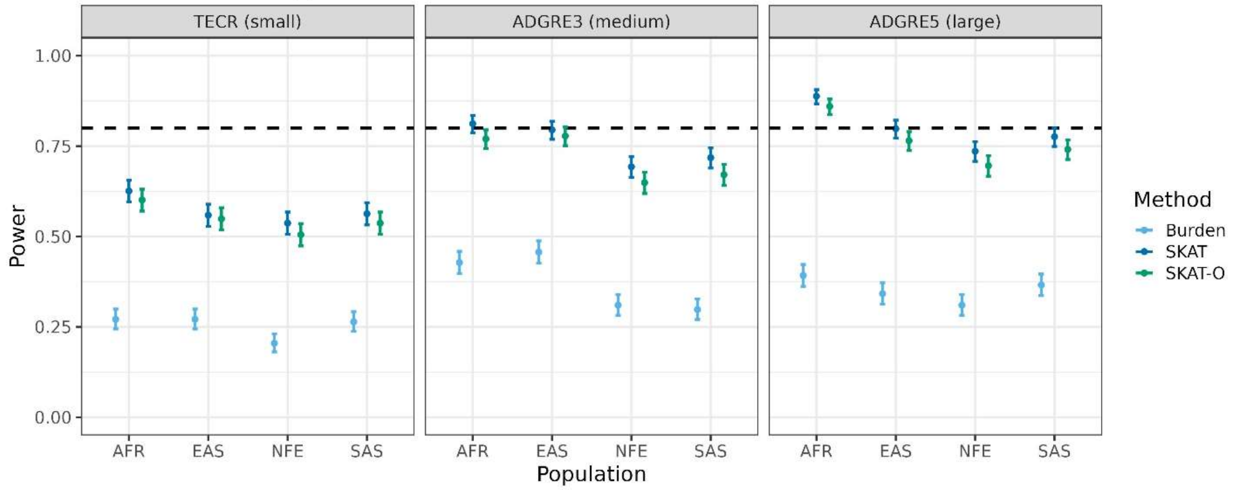


**Figure S10.** Power scenario results for 60% more causal functional variants than expected with $w_{fun,a} = 1.6$, $w_{fun,b} = 1$, and $w_{fun,c1} = w_{fun,c2} = 1$.
